# Controlling critical mistag-associated false discoveries in metagenetic data

**DOI:** 10.1101/2022.01.01.474695

**Authors:** Rodney T. Richardson

## Abstract

1. Metagenetic methods are commonplace within ecological and environmental research. One concern with these methods is the phenomenon of critical mistagging, where sequences from one sample are erroneously inferred to have originated from another sample due to errors in the attachment, PCR replication or sequencing of sample-specific dual-index tags. For studies using PCR-based library preparation on large sample sizes, the most cost-effective approach to limiting mistag-associated false detections involves using an unsaturated Latin square dual-indexing design. This allows researchers to estimate mistagging rates during sequencing but the statistical procedures for filtering out detections using this mistag rate have received little attention.
2. We propose a straightforward method to limit mistag-associated false discoveries during metabarcoding applications. We analyzed two Illumina metabarcoding datasets produced using unsaturated Latin square designs to explore the distribution of mistagged sequences across dual-index combinations on a per taxon basis. We tested these data for conformity to the assumptions that 1) mistagging follows a binomial distribution [i.e., *X* ~ *B*(*n, p*)] where *p*, the probability of a sequence being mistagged, varies minimally across taxa and 2) mistags are distributed uniformly across dual-index combinations. We provide R functions that estimate the 95^th^ percentile of expected mistags per dual-index combination for each taxon under these assumptions.
3. We show that mistagging rates were consistent across taxa within the datasets analyzed and that modelling mistagging as a binomial process with uniform distribution across dual-index combinations enabled robust control of mistag-associated false discoveries.
4. We propose that this method of taxon-specific filtering of detections based on the maximum mistags expected per dual-index combination should be broadly accepted during metagenetic analysis, provided that experimental and control sequence abundances per taxon are strongly correlated. When this assumption is violated, data may be better fit by assuming that the distribution of mistags across combinations follows Poisson characteristics [i.e., *X* ~ Pois(*λ*)], with *λ* empirically estimated from the abundance distribution of mistags among control samples. We provide a second R function for this case, though we have yet to observe such a dataset. Both functions and demonstrations associated with this work are freely available at https://github.com/RTRichar/ModellingCriticalMistags.

## Introduction

Metagenetic sequencing, often termed metabarcoding or amplicon sequencing when individual genetic markers are analyzed, is increasingly applicable to a variety of environmental and ecological research questions (Ficetola et al., 2008; Kartzinel et al., 2015; Keller et al., 2015; Willerslev et al., 2014). Part of the attractiveness of metagenetic sequencing is the capacity to analyze hundreds of samples per sequencing run through the use of sample dual-indexing during library preparation (e.g., Kozich et al. 2013). However, researchers quickly realized that the dual-indexing process introduces a form of sample cross-contamination into the resulting data where sequences from one sample are falsely inferred to have originated from another sample (Carlsen et al., 2012; Esling et al., 2015; Schnell et al., 2015). Artifactual sequences resulting from this process, known as ‘critical mistags’ or ‘tag jumps,’ generally represent a small fraction of the total sequences produced during metagenetic studies (Esling et al., 2015; Schnell et al., 2015). However, critical mistag rates can range widely depending on the library preparation methods used (Carøe & Bohmann, 2020) and thus mistagging represents an important issue to consider during the design and analysis of metagenetic studies.

Within the metagenetic literature, a number of techniques are used to append sample-specific indices during amplicon library preparation and these methods vary with regard to mistagging propensities, as reviewed in Bohmann et al. (2021). Critical mistagging can be nearly eliminated through the use of ligation-based library preparation procedures (Carøe & Bohmann, 2020) or by dual-indexing samples using an identity-based matrix of forward and reverse tags, referred to as ‘twin tagging’ (Schnell et al., 2015; Zepeda-Mendoza et al., 2016). Unfortunately, both of these solutions increase library preparation costs considerably, decreasing biological replication when funding is limited. A more cost-effective solution involves PCR-based indexing with an unsaturated Latin square design, where a fraction of dual-index combinations represent control samples used to measure the rate and pattern of critical mistagging within a sequencing dataset (Esling et al., 2015). Typically, these aspects of a sequencing run are measured using control samples we refer to as ‘no-library negatives,’ where specific dual-index combinations are left completely unused during library preparation. In the absence of alternate forms of contamination which obscure critical mistagging patterns, other forms of positive or negative controls can serve the purpose of quantifying mistagging rates, as outlined in Table 1. In our view, positive controls and no-library negative controls provide the greatest power for quantifying artifactual sequences in metagenetic studies. When positive control templates amplify with similar efficiency to experimental samples, relative contamination levels can be inferred across samples, since contaminants have to compete with positive control templates during amplification. For negative controls, no competitive templates are present and even small amounts of contamination will result in large numbers of contaminant sequences due to PCR amplification.

**Table 1:** Three types of control samples are commonly used in metabarcoding and metagenetic studies. What we refer to as ‘no-library negatives’ contain critical mistags only, while other forms of positive and negative controls can contain additional contaminant sequences.

| Control type | Origin and description | Artifacts detected |
| --- | --- | --- |
| Positive control | PCR products from a known control substrate that is amplifiable with the primers used on experimental sample substrates, ideally collected in the field using the same collection methods used on experimental samples | Critical mistags + lab and/or field contaminants |
| Negative control | PCR products from a known control substrate that is not amplifiable with the primers used on experimental sample substrates, ideally collected in the field using the same collection methods used on experimental samples | Critical mistags + lab and/or field contaminants |
| No-library negative control | No substrates are used for amplification and no library preparation reactions are conducted (i.e. no-library negatives represent dual-index combinations which are unused during library preparation) | Critical mistags |

Use of a Latin square design (Figure 1) minimizes mistagging by providing dual-index combinations that serve as mistag sinks for sample-containing mistag source combinations, decreasing the effective mistag rate among experimental samples (Esling et al., 2015). More importantly, this approach enables quantification of mistagging rates such that they may be statistically accounted for within research studies. While researchers have adopted this approach within the field, the methods for statistically controlling mistags are still unclear. This has led researchers to adopt a variety of approaches to limit critical mistag-associated discoveries (Cuff et al., 2021; Hänfling et al., 2016; Matesanz et al., 2019; McInnes et al., 2017; Zinger et al., 2021). To date, none of the current methods for filtering mistags are rooted in a statistical test which evaluates the probability that a given detection represents a critical mistag-associated false discovery.

**Figure 1:**
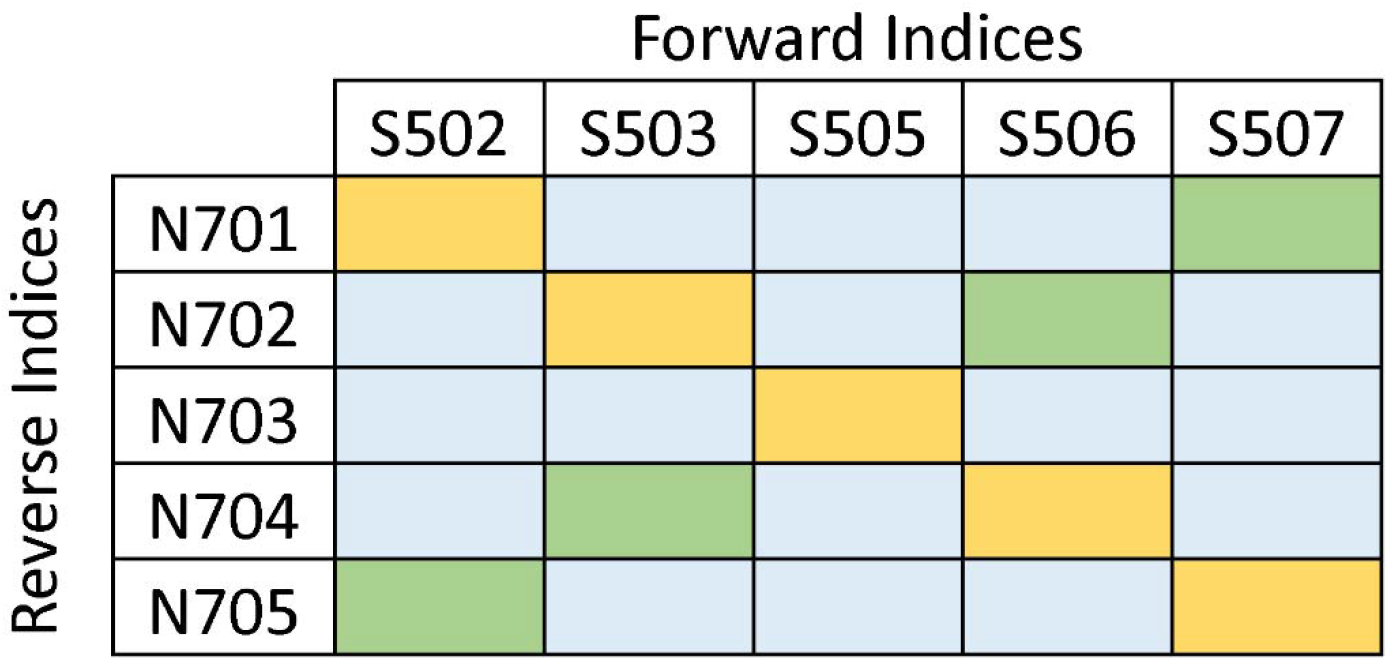
Illustration of a 36 percent unsaturated Latin square design (Esling et al., 2015) constructed for 16 experimental samples (blue dual-index combinations), 4 positive control samples (green dual-index combinations) and 5 no-library negative control samples (yellow dual-index combinations). When a distinct taxon not expected to be present among experimental samples is used for positive control samples, identification of contaminant sequences within positive controls in excess of critical mistag levels (i.e., levels observed among yellow combinations) allow for quantification of contamination levels incurred during sample processing. When total contaminant sequence abundances per taxon are strongly correlated between no-library negative controls and positive controls, field and/or lab contamination is likely rare and positive control contaminants predominately represent mistags.

Here, we propose a statistical framework for limiting experiment-wide mistag-associated false detections to a desired false discovery rate (FDR) within metagenetic studies. Such a method is important for both controlling false discoveries and maximizing sensitivity among metagenetic applications. Improving metagenetic sensitivity is particularly important when certain target taxa are expected to be rare or infrequently detected within samples, as is often the case for aquatic eDNA applications and metagenetic diet analysis studies (e.g. Cuff et al., 2021; Leese et al., 2021; Miya et al., 2015). We believe that the proposed methods will be useful for researchers wishing to conduct metagenetic applications cost-effectively and at large scales.

### Proposed method for limiting mistag-associated false discoveries

With the proposed method, any FDR limit between 0 and 1 can be specified and the following example used FDR of 0.05 since this is typical among research applications. The experiment-wide mistag-associated FDR can be limited to 0.05 through a series of steps, starting with the calculation of the 95^th^ percentile of expected mistags per dual-index combination. This involves calculating the upper 95 percent confidence limit of the mean number of mistags per dual-index combination (*M_cl_*) among control samples (Equation 1), where 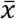 represents the mean number of mistags per dual-index combination, *σ* represents the standard deviation of the mean, *n* represent the number of control dual-index combinations and *t* is the *t*-statistic at the 0.975 quantile with *n* – 1 degrees of freedom. We then multiply *M_cl_* by the total number of dual-index combinations (*C*, equal to the total number of samples including controls) present in the dual-index matrix design to obtain the upper 95 percent confidence limit of the total number of experiment-wide mistags (*M_t_*, Equation 2). Total mistags can be divided by the total number of sequences (*S_t_*) to estimate the probability that any individual sequence was the result of a mistag event during sequencing (*P*, Equation 3).

We use *P* to estimate the expected distribution of experiment-wide mistags (*E_t_*) for each taxon in the dataset according to a binomial distribution where *A* represents the total number of sequences observed for the taxon (Equation 4, Figure 2A). The 95^th^ percentile of this distribution represents the 0.05 false discovery rate limit for the number of mistags expected per taxon (Figure 2A, dark blue line). We then account for the distribution of these mistags across individual dual-index combinations. When the relationship between total sequence abundance and mistag abundance per taxon is strong and no evidence of non-uniformity in mistag abundance is observed across the dual-index combinations, mistags are assumed to be distributed uniformly across 1 to *C* possible dual-index combinations (Equation 5, where *E_C_* is expected mistags per dual-index combination). We can simulate the uniform distribution of expected mistags across dual-index combinations *z* times, recording the maximum observed number of mistags per combination for each simulation, and estimate the 0.05 false discovery rate limit for the number of mistags per combination as the 95^th^ percentile of this distribution (Figure 2B, light blue line). Applying this to each taxon in a dataset, the relationship between total sequences, maximum expected mistags and maximum expected mistags per dual-index combination can be estimated (Figure 2C).

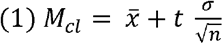

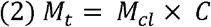

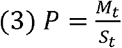

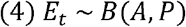

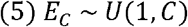

**Figure 2:**
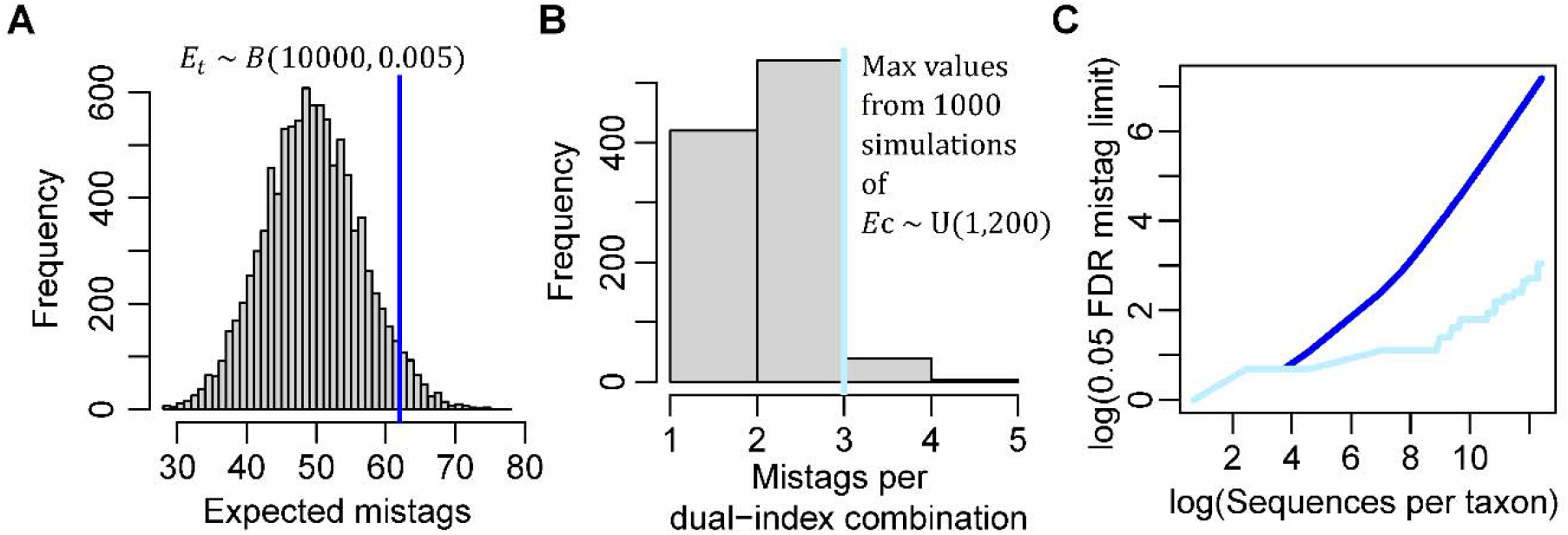
Illustration of the proposed method on a hypothetical sequencing dataset with a mistag probability of 0.005 and 200 dual-index sample combinations. For a taxon represented by 10,000 sequences across all samples, the binomial distribution is used to estimate the 95^th^ percentile of expected mistags (A, blue line). Through repeatedly simulating the uniform distribution of expected mistags across all samples, the 95^th^ percentile of expected mistags per dual-index combination can be calculated (B, light blue line). Repeating this approach for all taxa in the dataset, the relationship between the total sequence abundance, total mistags (blue line) and mistags per dual-index combination (light blue line) can be estimated at a false discovery rate of 0.05 (C).

The above example of evaluating the probability that a given discovery is the result of critical mistagging is provided with the assumption that mistags are distributed uniformly across all possible dual-index combinations. This assumption can be tested through bootstrapped comparison of the control sample mistag distribution to simulated uniform distributions using a Kolmogorov-Smirnov test as described below. When this assumption is violated, we provide an alternative R function where mistags are assumed to be distributed according to a Poisson process, with *λ* specifying the degree of non-uniformity and empirically estimated from the abundance distribution of mistags among control samples.

### Demonstration of the method on empirical data

The proposed method for limiting mistag-associated false discoveries was applied to two unpublished metabarcoding datasets produced using unsaturated Latin square designs, which allowed for an empirical test of the critical mistagging assumptions presented here. Dataset 1 consists of 30 bulk terrestrial arthropod community samples, 8 no-library negative control samples and 4 positive control samples (silver-haired bat, *Lasionycteris noctivagans*). Dataset 1 samples were amplified with BR2-BF2 primers (Elbrecht & Leese, 2017) using the three-step PCR-based library preparation methods of Richardson et al. (2019) and sequenced on an Illumina Nano flow cell (1 million reads, 500 cycles). Dataset 2 consists of 296 midgut DNA extracts from individual spiders and lady beetles and 7 no-library negative control samples. Dataset 2 samples were amplified with BR2-BF2 primers using the two-step PCR-based library preparation methods of Lange et al. (2014) and sequenced on a standard Illumina flow cell (15 million reads, 500 cycles). Both datasets were analyzed by merging forward and reverse reads using Pear (v.0.9.1; Zhang et al. (2014)) with a minimum trim length (-t) of 150 bp and a base call quality threshold (-q) of 20. Using VSEARCH (v.2.8.1; Rognes et al. (2016)), merged sequences were mapped against the arthropod COI dataset from Richardson et al. (2020), which was trimmed to the BR2-BF2 region using MetaCurator (v.1.0.1; Richardson et al. (2020)), with a query coverage threshold of 0.8 and an alignment percent identity threshold of 85. Data were summarized to the family level for downstream tests of the proposed method to ensure a high degree of classification sensitivity and accuracy across all taxa.

Within Dataset 1, sequence abundances per taxon were strongly correlated between no-library negative control and positive control samples (ordinary least squares regression on log-transformed abundances, *P* < 0.001, *R^2^* = 0.27), suggesting that critical mistags, not laboratory contamination, were the predominant source of arthropod sequences within positive control libraries. Thus, we included positive control sample data in calculating the experiment-wide mistag probability. A strong positive relationship between total mistag abundances and log-transformed total sequence abundances per taxon indicated minimal variation in mistagging propensity across taxa (Poisson GLM, *P* < 0.001, Nakagawa & Schielzeth Psuedo-*R^2^* = 0.94). The assumed uniform distribution of critical mistags across dual-indices was tested using a bootstrapped two-sample Kolmogorov-Smirnov test with 100 simulated uniform mistag distributions. Evidence for non-uniform distribution of mistags was weak and statistically insignificant (average *D* = 0.39, *P* > 0.1). The 95 percent confidence interval of mistags per dual-index combination ranged from 20 to 42, indicating an upper mistag probability of 0.005 for Dataset 1. Total mistag abundances were also strongly related to total sequence abundances per taxon in Dataset 2 (Poisson GLM, *P* < 0.001, Nakagawa & Schielzeth Psuedo-*R^2^* = 0.97). Marginal evidence for non-uniform distribution of mistagging events was found for Dataset 2 (average *D* = 0.73, *P* = 0.05), however, we continued our analysis assuming uniform distribution given the minimal number of control sample replicates (N = 7) and highly skewed abundance distribution of this diet analysis dataset.

The 95 percent confidence interval of mistags per dual-index combination ranged from 19 to 128, corresponding to an upper limit of 0.003 with regard to the Dataset 2 mistagging probability.

False discovery rate thresholds visually appeared well-fit to the data, where all instances of mistags per taxon and nearly all instances of mistags per taxon per dual-index combination were below the 0.05 false discovery rate limit for both datasets (Figure 3). The proposed method was quantitatively evaluated by estimating the experiment-wide false discovery rate among control dual-index combinations according to Equation 6, where *F* represents the number of control sample detections exceeding the false discovery rate threshold, *N* represents the number of control samples and *T* represents the number of taxa present in the study. Importantly, *T* included only taxa represented by at least 2,000 sequences to ensure that false discovery rate estimates were calculated from abundant taxa with reasonable likelihood of producing mistag-associated false discoveries. Dataset 1 exhibited a false discovery rate of 0.013, which was significantly less than 0.05 (Pearson’s *X^2^* test with Yates’ continuity correction, *P* < 0.001, *X^2^* = 10.871, 95% CI: 0.005 to 0.0309). Dataset 2 yielded a false discovery rate of 0.026, which was not significantly different from 0.05 (Pearson’s *X^2^* test with Yates’ continuity correction, *P* = 0.480, *X^2^* = 0.498, 95% CI: 0.0045 to 0.0993). With regard to metagenetic sensitivity, the proposed method resulted in the removal of 34 percent of non-control detections from Dataset 1 and 46 percent of detections from Dataset 2. No taxa were removed during mistag filtering of either dataset. Unless a considerably larger number of sequences are present in control samples relative to experimental samples for a given taxon (i.e., a contamination event), this method would not be expected to reduce experiment-wide taxonomic richness.

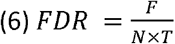

**Figure 3:**
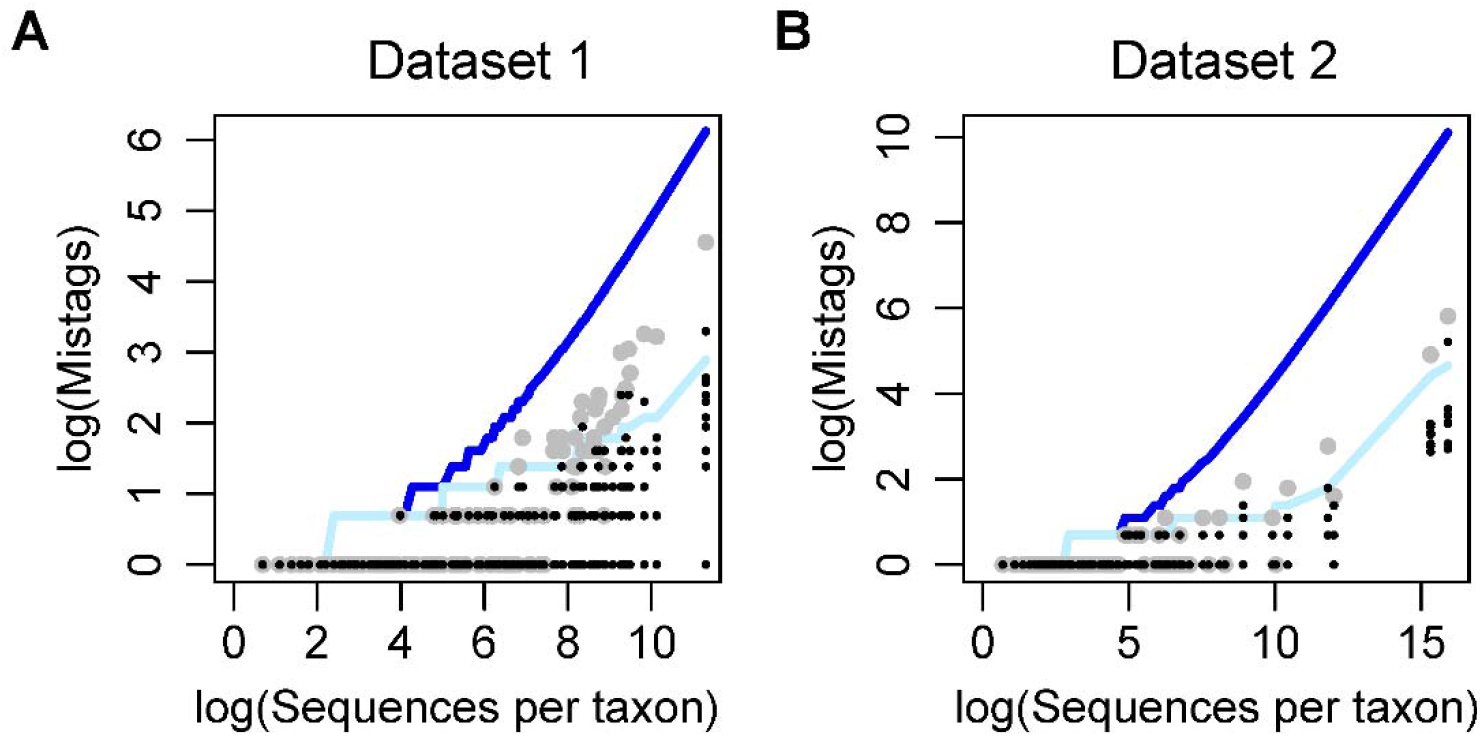
Expected vs. observed assessment of the proposed mistag control methods on Dataset 1 (A) and Dataset 2 (B). For all plots, the dark blue line indicates the maximum expected number of mistagged sequences per taxon across the whole dataset at a false discovery rate of 0.05. Light blue lines indicate the maximum expected mistags per taxon per dual-index combination at a false discovery rate of 0.05, assuming uniform distribution of mistags across combinations. Large grey points show the observed experiment-wide number of mistags per taxon and small black points show the observed number of mistags per taxon per dual-index combination among positive and negative control samples. A pseudocount of 1 was added to the x and y-axis to facilitate log-transformation of zeros for both plots. Expected mistags per taxon per dual-index combination at a false discovery rate of 0.05 were estimated using 25,000 simulations.

With regard to detection sensitivity, varying the FDR limit from (1-1×10^-100^) to 5×10^-7^ resulted in an exponential decay of the proportion of inferred true positive detections toward asymptotes of approximately 0.63 for Dataset 1 and 0.51 for Dataset 2 (Figure 4). Typical FDR limits between 0.2 and 0.01 resulted in observed sensitivities ranging from 0.73 to 0.68 for Dataset 1 and 0.61 to 0.53 for Dataset 2.

**Figure 4:**
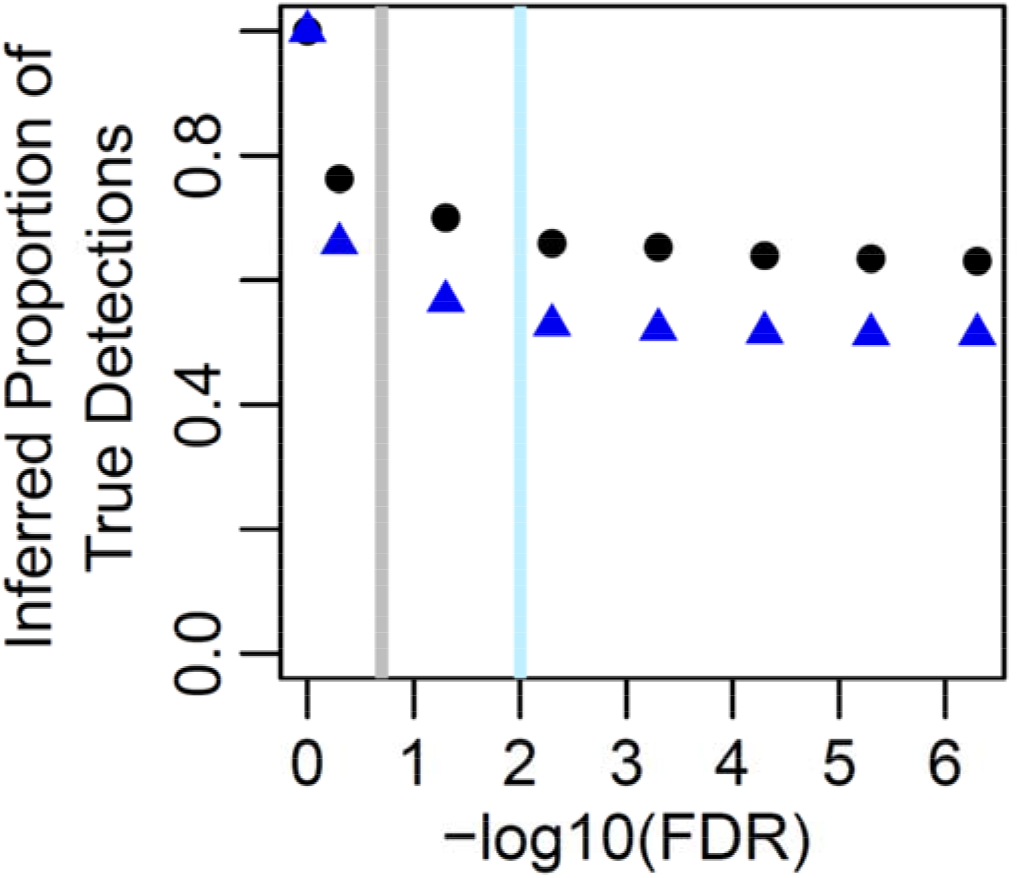
Approximate relationship between the detection confidence limit, FDR (x-axis), and detection sensitivity, the proportion of original detections inferred to be true positives (y-axis). Results are shown for Dataset 1 (black circles) and Dataset 2 (blue triangles). The range of confidence levels typically within research contexts are emphasized with a grey line (FDR = 0.2, 0.8 confidence) and light blue line (FDR = 0.01, 0.99 confidence).

## Concluding remarks

We provide a statistical framework for evaluating the probability that a given metagenetic identification is the result of critical mistagging for experiments that include an unsaturated Latin square dual-indexing design. The method relies upon the accurate estimation of the proportion of mistagged sequences, *P*, for each sequencing dataset within an experiment. Smaller numbers of control samples will result in greater uncertainty in *P*. Since we recommend using the upper limit of the 95 percent confidence interval of *P*, the estimated probability that a detection is the result of mistagging will likely be inflated when few control sample are used, lowering sensitivity for true detections and increasing the false negative rate of the experiment. Thus, researchers should carefully consider positive and negative control sample sizes to ensure sufficient power for the precise estimation of mistagging probabilities in order to maximize detection power. To this end, probabilistic removal of potential mistag-associated false positives will inevitably generate false negative results. For the datasets analyzed, a typical FDR threshold of 0.05 resulted in the removal of 30 to 43 percent of total detections (Figure 4). Since we do not know the fraction of true positive relative to false positive inferences, we cannot precisely estimate the rate of false negatives resulting from this process. Regardless, the proposed method is not be expected to completely remove a taxonomic unit from any dataset unless that taxonomic unit exhibits much greater sequence abundance in control samples relative to experimental samples, a situation indicative of a contamination event. Thus, experiment-wide sensitivity for any specific taxonomic unit is expected to approximately 1 using the proposed method.

Analyses of the data presented here were conducted at the family level. Given the sequence classification methods and reference database used, we felt this was appropriate to ensure a high degree of bioinformatic precision for the demonstrations provided. Researchers wishing to use this method during analyses with low classification precision at the desired taxonomic rank, such as genus or species for most systems, may wish to rely on sequence cluster units as opposed to classifications of individual sequence cases. Sequence clustering allows for more precise handling of the sequence variants present within metagenetic datasets (Callahan et al., 2016; Zou et al., 2020), though the relationships between these types of sequence clusters and taxonomic affiliations can be uncertain in some cases (Edgar, 2018; Schloss, 2021).

Multiple methods have been proposed for limiting mistag-associated false discoveries in metagenetic data, but most require considerable increases in library preparation costs (Carøe & Bohmann, 2020; Schnell et al., 2015; Zepeda-Mendoza et al., 2016). PCR-based library preparation with a Latin square design represents the most cost-effective approach to date, but precise statistical methods for evaluating mistag-associated false discovery rates have not been formulated. The proposed method is based on mistagging patterns observed across multiple datasets and appears to provide for robust control of mistag-associated false discoveries. It is provided in an easy to use format to facilitate uptake by end-users and further confirmations of the method on a more diverse array of datasets is encouraged. We believe this method will be useful in a wide variety of metagenetics applications, though it is important to note that some applications require detection confidence levels that may exceed what can be provided by this method. In these cases, more costly library preparation protocols or more species-specific molecular detection methods may be warranted. Current implementations of metagenetics techniques are highly variable and continued efforts are needed to quantify trade-offs between detection confidence, taxonomic breadth of detection and cost in different research contexts. Improved inference of mistag-associated false discoveries will aid in these efforts.

## Data Accessibility Statement

Demonstrations of the proposed method on simulated data can be found at https://github.com/RTRichar/ModellingCriticalMistags.

## References

Bohmann, K., Elbrecht, V., Carøe, C., Bista, I., Leese, F., Bunce, M., Yu, D. W., Seymour, M., Dumbrell, A., & Creer, S. (2021). Strategies for sample labelling and library preparation in DNA metabarcoding studies. https://doi.org/10.22541/au.162141261.10649593/v1

Callahan, B. J., McMurdie, P. J., Rosen, M. J., Han, A. W., Johnson, A. J. A., & Holmes, S. P. (2016). DADA2: High-resolution sample inference from Illumina amplicon data. Nature Methods, 13(7), 581–583. https://doi.org/10.1038/nmeth.3869

Carlsen, T., Aas, A. B., Lindner, D., Vrålstad, T., Schumacher, T., & Kauserud, H. (2012). Don’t make a mista(g)ke: Is tag switching an overlooked source of error in amplicon pyrosequencing studies? Fungal Ecology, 5(6), 747–749. https://doi.org/10.1016/j.funeco.2012.06.003

Carøe, C., & Bohmann, K. (2020). Tagsteady: A metabarcoding library preparation protocol to avoid false assignment of sequences to samples. Molecular Ecology Resources, 20(6), 1620–1631. https://doi.org/10.1111/1755-0998.13227

Cuff, J. P., Drake, L. E., Tercel, M. P. T. G., Stockdale, J. E., Orozco-terWengel, P., Bell, J. R., Vaughan, I. P., Müller, C. T., & Symondson, W. O. C. (2021). Money spider dietary choice in pre- and post-harvest cereal crops using metabarcoding. Ecological Entomology, 46(2), 249–261. https://doi.org/10.1111/een.12957

Edgar, R. C. (2018). Updating the 97% identity threshold for 16S ribosomal RNA OTUs. Bioinformatics, 34(14), 2371–2375. https://doi.org/10.1093/bioinformatics/bty113

Elbrecht, V., & Leese, F. (2017). Validation and development of COI metabarcoding primers for freshwater macroinvertebrate bioassessment. Frontiers in Environmental Science, 5, 11. https://doi.org/10.3389/fenvs.2017.00011

Esling, P., Lejzerowicz, F., & Pawlowski, J. (2015). Accurate multiplexing and filtering for high-throughput amplicon-sequencing. Nucleic Acids Research, 43(5), 2513–2524. https://doi.org/10.1093/nar/gkv107

Ficetola, G. F., Miaud, C., Pompanon, F., & Taberlet, P. (2008). Species detection using environmental DNA from water samples. Biology Letters, 4(4), 423–425. https://doi.org/10.1098/rsbl.2008.0118

Hänfling, B., Lawson Handley, L., Read, D. S., Hahn, C., Li, J., Nichols, P., Blackman, R. C., Oliver, A., & Winfield, I. J. (2016). Environmental DNA metabarcoding of lake fish communities reflects long-term data from established survey methods. Molecular Ecology, 25(13), 3101–3119. https://doi.org/10.1111/mec.13660

Kartzinel, T. R., Chen, P. A., Coverdale, T. C., Erickson, D. L., Kress, W. J., Kuzmina, M. L., Rubenstein, D. I., Wang, W., & Pringle, R. M. (2015). DNA metabarcoding illuminates dietary niche partitioning by African large herbivores. Proceedings of the National Academy of Sciences, 112(26), 8019. https://doi.org/10.1073/pnas.1503283112

Keller, A., Danner, N., Grimmer, G., Ankenbrand, M., von der Ohe, K., von der Ohe, W., Rost, S., Härtel, S., & Steffan-Dewenter, I. (2015). Evaluating multiplexed next-generation sequencing as a method in palynology for mixed pollen samples. Plant Biology, 17(2), 558–566. https://doi.org/10.1111/plb.12251

Kozich, J. J., Westcott, S. L., Baxter, N. T., Highlander, S. K., & Schloss, P. D. (2013). Development of a dual-index sequencing strategy and curation pipeline for analyzing amplicon sequence data on the MiSeq Illumina sequencing platform. Applied and Environmental Microbiology, 79(17), 5112–5120. PubMed. https://doi.org/10.1128/AEM.01043-13

Lange, V., Böhme, I., Hofmann, J., Lang, K., Sauter, J., Schöne, B., Paul, P., Albrecht, V., Andreas, J. M., Baier, D. M., Nething, J., Ehninger, U., Schwarzelt, C., Pingel, J., Ehninger, G., & Schmidt, A. H. (2014). Cost-efficient high-throughput HLA typing by MiSeq amplicon sequencing. BMC Genomics, 15(1), 63. https://doi.org/10.1186/1471-2164-15-63

Leese, F., Sander, M., Buchner, D., Elbrecht, V., Haase, P., & Zizka, V. M. A. (2021). Improved freshwater macroinvertebrate detection from environmental DNA through minimized nontarget amplification. Environmental DNA, 3(1), 261–276. https://doi.org/10.1002/edn3.177

Matesanz, S., Pescador, D. S., Pías, B., Sánchez, A. M., Chacón-Labella, J., Illuminati, A., de la Cruz, M., López-Angulo, J., Marí-Mena, N., Vizcaíno, A., & Escudero, A. (2019). Estimating belowground plant abundance with DNA metabarcoding. Molecular Ecology Resources, 19(5), 1265–1277. https://doi.org/10.1111/1755-0998.13049

McInnes, J. C., Alderman, R., Deagle, B. E., Lea, M.-A., Raymond, B., & Jarman, S. N. (2017). Optimised scat collection protocols for dietary DNA metabarcoding in vertebrates. Methods in Ecology and Evolution, 8(2), 192–202. https://doi.org/10.1111/2041-210X.12677

Miya, M., Sato, Y., Fukunaga, T., Sado, T., Poulsen, J. Y., Sato, K., Minamoto, T., Yamamoto, S., Yamanaka, H., Araki, H., Kondoh, M., & Iwasaki, W. (2015). MiFish, a set of universal PCR primers for metabarcoding environmental DNA from fishes: Detection of more than 230 subtropical marine species. Royal Society Open Science, 2(7), 150088. https://doi.org/10.1098/rsos.150088

Richardson, R. T., Curtis, H. R., Matcham, E. G., Lin, C.-H., Suresh, S., Sponsler, D. B., Hearon, L. E., & Johnson, R. M. (2019). Quantitative multi-locus metabarcoding and waggle dance interpretation reveal honey bee spring foraging patterns in Midwest agroecosystems. Molecular Ecology, 28(3), 686–697. https://doi.org/10.1111/mec.14975

Richardson, R. T., Sponsler, D. B., McMinn-Sauder, H., & Johnson, R. M. (2020). MetaCurator: A hidden Markov model-based toolkit for extracting and curating sequences from taxonomically-informative genetic markers. Methods in Ecology and Evolution, 11(1), 181–186. https://doi.org/10.1111/2041-210X.13314

Rognes, T., Flouri, T., Nichols, B., Quince, C., & Mahé, F. (2016). VSEARCH: A versatile open source tool for metagenomics. PeerJ, 4, e2584. https://doi.org/10.7717/peerj.2584

Schloss, P. D. (2021). Amplicon Sequence Variants Artificially Split Bacterial Genomes into Separate Clusters. MSphere, 6(4), e0019121. https://doi.org/10.1128/mSphere.00191-21

Schnell, I. B., Bohmann, K., & Gilbert, M. T. P. (2015). Tag jumps illuminated – reducing sequence-to-sample misidentifications in metabarcoding studies. Molecular Ecology Resources, 15(6), 1289–1303. https://doi.org/10.1111/1755-0998.12402

Willerslev, E., Davison, J., Moora, M., Zobel, M., Coissac, E., Edwards, M. E., Lorenzen, E. D., Vestergård, M., Gussarova, G., Haile, J., Craine, J., Gielly, L., Boessenkool, S., Epp, L. S., Pearman, P. B., Cheddadi, R., Murray, D., Bråthen, K. A., Yoccoz, N., & Taberlet, P. (2014). Fifty thousand years of Arctic vegetation and megafaunal diet. Nature, 506(7486), 47–51. https://doi.org/10.1038/nature12921

Zepeda-Mendoza, M. L., Bohmann, K., Carmona Baez, A., & Gilbert, M. T. P. (2016). DAMe: A toolkit for the initial processing of datasets with PCR replicates of double-tagged amplicons for DNA metabarcoding analyses. BMC Research Notes, 9(1), 255. https://doi.org/10.1186/s13104-016-2064-9

Zhang, J., Kobert, K., Flouri, T., & Stamatakis, A. (2014). PEAR: A fast and accurate Illumina Paired-End reAd mergeR. Bioinformatics, 30(5), 614–620. https://doi.org/10.1093/bioinformatics/btt593

Zinger, L., Lionnet, C., Benoiston, A.-S., Donald, J., Mercier, C., & Boyer, F. (2021). MetabaR: An r package for the evaluation and improvement of DNA metabarcoding data quality. Methods in Ecology and Evolution, 12(4), 586–592. https://doi.org/10.1111/2041-210X.13552

Zou, Q., Lin, G., Jiang, X., Liu, X., & Zeng, X. (2020). Sequence clustering in bioinformatics: An empirical study. Briefings in Bioinformatics, 21(1), 1–10. https://doi.org/10.1093/bib/bby090

